# Microinjection, gene knockdown, and CRISPR-mediated gene knock-in in the hard coral, *Astrangia poculata*

**DOI:** 10.1101/2023.11.16.567385

**Authors:** Jacob F. Warner, Ryan Besemer, Alicia Schickle, Erin Borbee, Isabella V. Changsut, Koty Sharp, Leslie S. Babonis

## Abstract

Cnidarians have become valuable models for understanding many aspects of developmental biology including the evolution of body plan diversity, novel cell type specification, and regeneration. Most of our understanding of gene function during early development in cnidarians comes from a small number of experimental systems including the sea anemone, *Nematostella vectensis*. Few molecular tools have been developed for use in hard corals, limiting our understanding of this diverse and ecologically important clade. Here, we report the development of a suite of tools for manipulating and analyzing gene expression during early development in the northern star coral, *Astrangia poculata*. We present methods for gene knockdown using short hairpin RNAs, gene overexpression using exogenous mRNAs, and endogenous gene tagging using CRISPR-mediated gene knock-in. Combined with our ability to control spawning in the laboratory, these tools make *A. poculata* a tractable experimental system for investigative studies of coral development. Further application of these tools will enable functional analyses of embryonic patterning and morphogenesis across Anthozoa and open new frontiers in coral biology research.

**Summary Statement:** This study reports the development of the first transgenic knock-in coral, providing the opportunity to track the behavior of various cell types during early coral development.

## Introduction

Recent advances in the techniques available for genetic manipulation have enabled the ability to perturb and analyze gene function in a broad range of animal phyla (Crawford et al., 2020; Oulhen et al., 2022; Presnell and Browne, 2021; Tinoco et al., 2023). These advancements make it possible to interrogate the evolution of development in taxa representing extreme variation in animal body plans. Among cnidarians, representatives of both Anthozoa (corals, sea anemones, etc) and Medusozoa (hydroids, jellyfish, etc) have emerged as highly tractable experimental systems; however, our understanding of development in these clades arises from investigation of only few species. As an example, the molecular regulation of embryogenesis appears to be well-studied in Anthozoa, yet most of our conclusions about this large and diverse clade of cnidarians derive from studies of the starlet sea anemone, *Nematostella vectensis* (Layden et al., 2016). Investigative studies of gene function in other anthozoans have been challenged by lack of accessibility to gametes, protected status of the adult, and a dearth of molecular tools.

The northern star coral, *Astrangia poculata*, is an attractive experimental organism for research on hard coral (scleractinian) development. This facultatively symbiotic, gonochoristic coral is found in high abundance in coastal waterways from the southern Caribbean to Cape Cod MA, USA (Dimond et al., 2013) and is listed by the IUCN as a species of “least concern”. In the late summer, when gametogenesis is at its peak in *A. poculata*, colonies can be collected from near-shore locations, induced to spawn *ex situ*, and their hardy, transparent larvae can be conveniently reared in laboratory conditions (Szmant-Froelich et al., 1980). The considerable ease of access to coral colonies combined with the ability to precisely control the timing of fertilization in the laboratory provides the opportunity to genetically manipulate early-stage embryos. Here, we describe the development of molecular tools for investigating gene function during early development in *A. poculata*. These tools establish *A. poculata* as a tractable research organism for functional studies in corals and, more broadly, as a viable system for comparative studies of cnidarian evolution and development

## Results and Discussion

### Spawning and microinjection of *Astrangia poculata*

To establish methods for manipulating gene function during embryogenesis in *A. poculata*, we collected wild adult colonies and induced spawning at precise times by raising the water temperature quickly from 19.5°C to 27-28°C in benchtop containers. Gamete release began 1-1.5 hours after heating (**Fig 1A,B**). To perform microinjection, we concentrated fertilized zygotes in a small volume of seawater and pipetted them gently onto the top of a piece of 100μm nylon mesh secured with modeling clay to the bottom of a 35 mm petri dish filled with filtered sea water (**Fig 1C,D**). The nylon mesh serves to cradle the individual zygotes during microinjection and the clay ensures the mesh can be removed easily to facilitate recovery of injected zygotes. At room temperature (∼22°C), first cleavage occurs after approximately 90 minutes, allowing for injection of a large number of zygotes. First cleavage in *A. poculata* is holoblastic, resulting in complete segregation of the first two embryonic cells. We demonstrated this by injecting two different dyes at the 2-cell stage and observing conserved segregation of the dyes later in development (**Fig 1E)**. By contrast, in *N. vectensis* complete segregation of embryonic cells is not observed until the 8-cell stage (Fritzenwanker et al., 2007). Thus, single-cell injections at the 2-cell stage in *A. poculata* could be used to knockdown or overexpress gene products in half of the embryo, facilitating studies of cell-cell communication during early development. To test the feasibility of using exogenous mRNAs for over- and mis-expression assays in *A. poculata*, we injected mRNA encoding a transcription factor (T-cell factor/TCF) isolated from *N. vectensis*, fused to a fluorescent protein (NvTCF-venus) (Röttinger et al., 2012). Blastula stage embryos exhibited nuclear localization of the fluorescent fusion protein as anticipated, and expression was maintained in healthy, dividing cells throughout early development (**Fig 1F**). These results demonstrate robust expression of exogenous transgenes, opening the possibility of using mis-expression approaches to study gene regulatory network diversification across species.

**Fig. 1.**
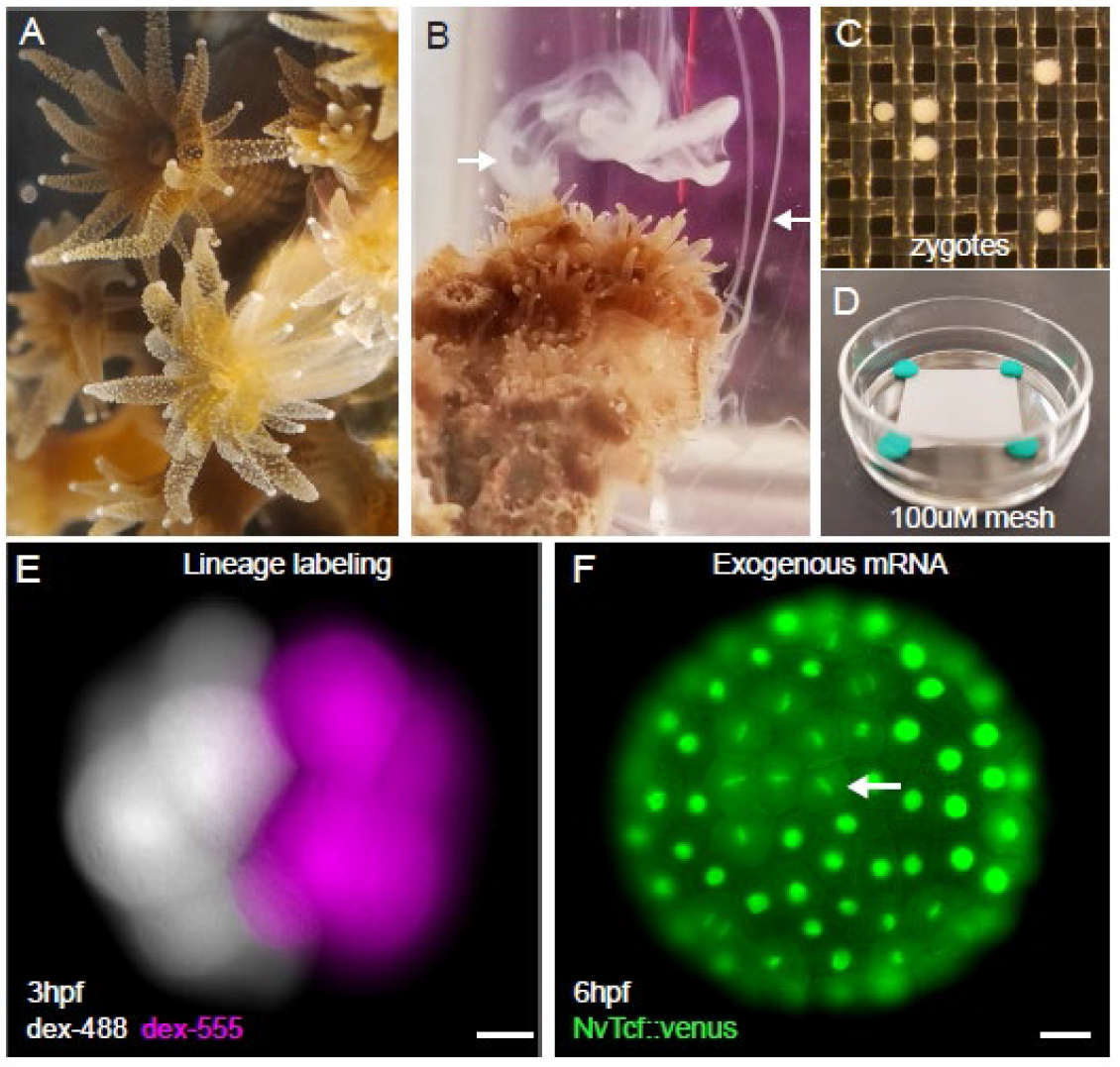
*Astrangia poculata* is a tractable research organism for functional molecular studies in corals. (A) *A. poculata* colony showing extended polyps. (B) *A. poculata* colony during spawning; arrows point to sperm emerging from two polyps. (C,D) Zygotes resting on nylon mesh fixed in the bottom of a 35mm petri dish in preparation for microinjection. (E) Live image of a 16-cell stage embryo injected at the 2-cell stage with two different dextran dyes (dex-488, dex-555). (F) Injection of mRNA encoding a TCF-venus fusion protein from *Nematostella vectensis* (NvTCF::venus) demonstrates proper translation and nuclear localization of exogenous mRNAs; arrow points to cells in M-phase. Scale bars = 20μm.

### Short hairpin RNAs enable efficient gene knockdown

RNA interference techniques have become indispensable for studies of early development as they allow for efficient, robust knockdown of gene function across cell and tissue types. Among cnidarians, RNAi technologies have been successful for manipulating gene function in a wide array of taxa, including both hydrozoans (DuBuc et al., 2020; Lohmann et al., 1999; Masuda-Ozawa et al., 2022; Quiroga-Artigas et al., 2020) and anthozoans (Dunn et al., 2007; He et al., 2018; Yuyama et al., 2021). Recently, short hairpin RNAs (shRNA) have become a widely adopted approach for RNA interference as they can be synthesized in the lab, thereby enabling cost-effective silencing of numerous target genes (He et al., 2018). We tested the efficacy of shRNA knockdown in *A. poculata* by inhibiting the activity of Fibroblast growth factor A1 (FgfA1). In *N. vectensis*, the role of FGF signaling during embryonic patterning has been well-studied (Gilbert et al., 2022; Rentzsch et al., 2008; Sinigaglia et al., 2015) and this pathway is known to be required for the formation of the apical tuft, a sensory structure at the aboral end of the larva from which a group of long cilia emerge (Rentzsch et al., 2008). While the apical tuft is found throughout sea anemones, most coral larvae lack this structure. The presence of an apical tuft in the larval stage of *A. poculata* (Szmant-Froelich et al., 1980) provides an opportunity to investigate the developmental mechanisms driving the convergent evolution of this structure across anthozoans.

To inhibit Fgf1A function in *A. poculata*, we injected zygotes with either a shRNA targeting the 3’ end of the FgfA1 transcript (see Materials and Methods) or a scrambled control shRNA (Karabulut et al., 2019). Animals were then raised to the larval stage at room temperature and inspected for evidence of an apical tuft. At 48 hours post fertilization (hpf), 10/10 of the knockdown larvae lacked apical tuft cilia, consistent with an inhibition of FgfA1 function (**Fig 2A**). To further confirm the role of FgfA1 in regulating apical tuft development, we treated a separate group of zygotes with the MEK/ERK inhibitor SU5402 (20μM), which has previously been shown to inhibit FgfA1-mediated control of apical tuft development in *N. vectensis* (Rentzsch et al., 2008). Treatment with SU5402 effectively phenocopied FgfA1 shRNA knockdown (**Fig 2B**), resulting in the complete loss of an apical tuft in 12/12 larvae at 48hpf.

**Fig. 2.**
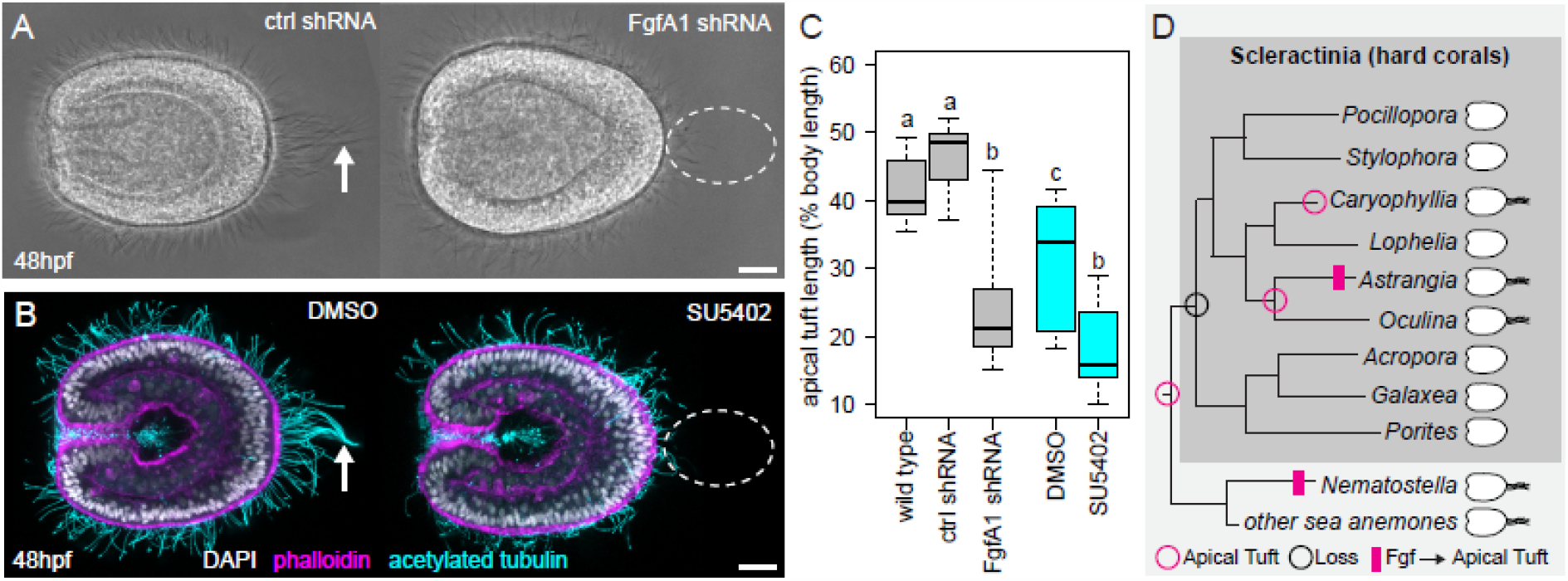
Knockdown of FgfA1 induces loss of the apical tuft. (A) Live images of 48 hpf larvae injected with scrambled control shRNA (ctrl shRNA) or FgfA1 shRNA. (B) Images of fixed, 48 hpf larvae treated with 20μM SU5402 or vehicle control (DMSO) and stained with DAPI (nuclei), phalloidin (F-actin), and anti-acetylated tubulin antibody (cilia). Arrows in A,B point to apical tuft cilia and dotted circles indicate loss of apical tuft. The oral pole is to the left in A,B; scale bars = 20μm. (C) Quantitative analysis of apical tuft cilia length in the shRNA experiment (grey boxes) and pharmacological experiment (cyan boxes). Box plots are presented as: median – middle line, 25^th^ and 75^th^ percentiles – box, 5^th^ and 95^th^ percentiles – whiskers. Sample sizes for each treatment: wild type N=10, ctrl shRNA N=8, FgfA1 shRNA N=10, DMSO N=10, SU5402 N=12. P-values from ANOVA with TukeyHSD posthoc: wild type vs. ctrl shRNA: p=0.5839021, wild type vs FgfA1 shRNA: p=0.0000064, ctrl shRNA vs FgfA1 shRNA: p= 0.0000001, wild type vs DMSO: p= 0.0154365, DMSO vs SU5402: p= 0.0004114, FgfA1 shRNA vs SU5402: p= 0.3147784. Letters indicate groups that are significantly different. (D) Cladogram of hard corals and sea anemones plotting the distribution of taxa with a larval apical tuft (cartoons, right). The apical tuft was likely lost in the ancestor of Scleractinia (black circle) and regained in the ancestor of the clade containing *Astrangia* and *Oculina* and at least one species of Caryophyllia (magenta circles). An Fgf signaling pathway controls apical tuft development in *Astrangia poculata* (this study) and *Nematostella vectensis* (Rentzsch et al., 2008). The cladogram was inferred from two studies of overlapping taxa (Kitahara et al., 2010; McFadden et al., 2021). References indicating presence/absence of apical tuft by taxon: *Pocillopora* (Tran and Hadfield, 2013), *Stylophora* (Atoda, 1951), *Caryophyllia* (Tranter et al., 1982), *Lophelia* (Larsson et al., 2014), *Astrangia* (Szmant-Froelich et al., 1980), *Oculina* (Brooke and Young, 2003), *Acropora* (Hayward et al., 2015), *Galaxea* (Atoda, 1951), *Porites* (Santiago-Valentín et al., 2022), *Nematostella* (Hand and Uhlinger, 1992), other sea anemones: *Anthopleura* (Chia and Koss, 1979), *Exaiptasia* (Bucher et al., 2016), *Gonactinia* (Chia et al., 1989).

These phenotypes were quantified by measuring the length of the longest cilium at the aboral end of each larva. The aboral cilia of the FgfA1 knockdown animals were significantly shorter than those in both the wild type and control shRNA-injected larvae (**Fig 2C**). Likewise, we observed a significant reduction in the length of cilia at the aboral pole in SU5402-treated larvae, relative to DMSO controls (Fig 2C). These experiments demonstrate that shRNA injection is a robust method for gene knockdown in corals and confirm that apical tuft development requires similar signaling pathways in two distantly related anthozoans (*N. vectensis* and *A. poculata*) **(Fig 2D**). Pharmacological inhibition of FGF signaling has also been shown to inhibit settlement and metamorphosis in *Acropora millepora*, a species that lacks an apical tuft (Cleves et al., 2018; Strader et al., 2018). With access to this inexpensive and robust method for gene knockdown it is now possible to interrogate the evolution of the FGF signaling pathway controlling apical sensory organ development in cnidarians with diverse larval body plans.

### Development of a transgenic knock-in coral to study cnidocyte development

Genome editing approaches using CRISPR/Cas9 technology have already been used for loss-of-function analysis in a variety of cnidarians including the sea anemone *Nematostella vectensis*, the hard coral *Acropora millepora*, and the hydroids *Hydractinia symbiolongicarpus* and *Clytia hemisphaerica* (Cleves et al., 2018; Gahan et al., 2017; Ikmi et al., 2014; Momose et al., 2018). Endogenous tagging of native proteins with fluorescent markers using CRISPR-mediated homology-directed repair (HDR) has further enabled precise tagging of individual proteins and careful analysis of protein activity in vivo in *N. vectensis* and *H. symbiolongicarpus* (Ikmi et al., 2014; Paix et al., 2023; Sanders et al., 2018). To date, however, successful gene knock-in in corals has not been reported. To establish a method for CRISPR-mediated gene knock-in in *A. poculata*, we tested a method that uses PCR-generated micro-homology fragments to induce HDR after CRISPR-Cas9 cleavage (Seleit et al., 2021). The benefit of this method is that knock-in repair templates can be constructed rapidly and inexpensively by PCR, without the need for cloning. To test the efficacy of CRISPR-mediated knock-in, we tagged the cnidocyte-specific marker gene, Minicollagen3 (*Mcol3*), with the fluorescent protein, mNeonGreen (mNeon). Minicollagens are found only in cnidocytes, making the expression of *Mcol3* a specific and robust marker of cnidocyte development (David et al., 2008).

We designed two single guide RNAs (sgRNAs) targeting the stop codon of the last exon of *Mcol3* and used an HDR repair template to insert the sequence of mNeon downstream of and in frame with *Mcol3* (**Fig 3A**). After injecting this repair template along with the sgRNAs and Cas9 protein, we observed positive fluorescent signal in developing cnidocytes beginning at 36hpf in approximately 10% (8/80) of injected larvae (**Fig 3B**). Most of the knock-in larvae (7/8) exhibited mosaic expression of Mcol3::mNeon, a common outcome of CRISPR-mediated genome editing likely representing a repair event that occurred at later embryonic stages. We confirmed positive integration using PCR with primers that flank mNeon to discriminate wild type alleles from mutant alleles with gel electrophoresis (**Fig 3C**). Using *in situ* hybridization, we confirmed that the knock-in construct recapitulated endogenous expression, showing that *Mcol3* is expressed in a salt and pepper pattern in the ectoderm during embryogenesis in *A. poculata* (**Fig 3D**), a pattern consistent with the development of cnidocytes in *N. vectensis* (Zenkert et al., 2011).

**Fig. 3.**
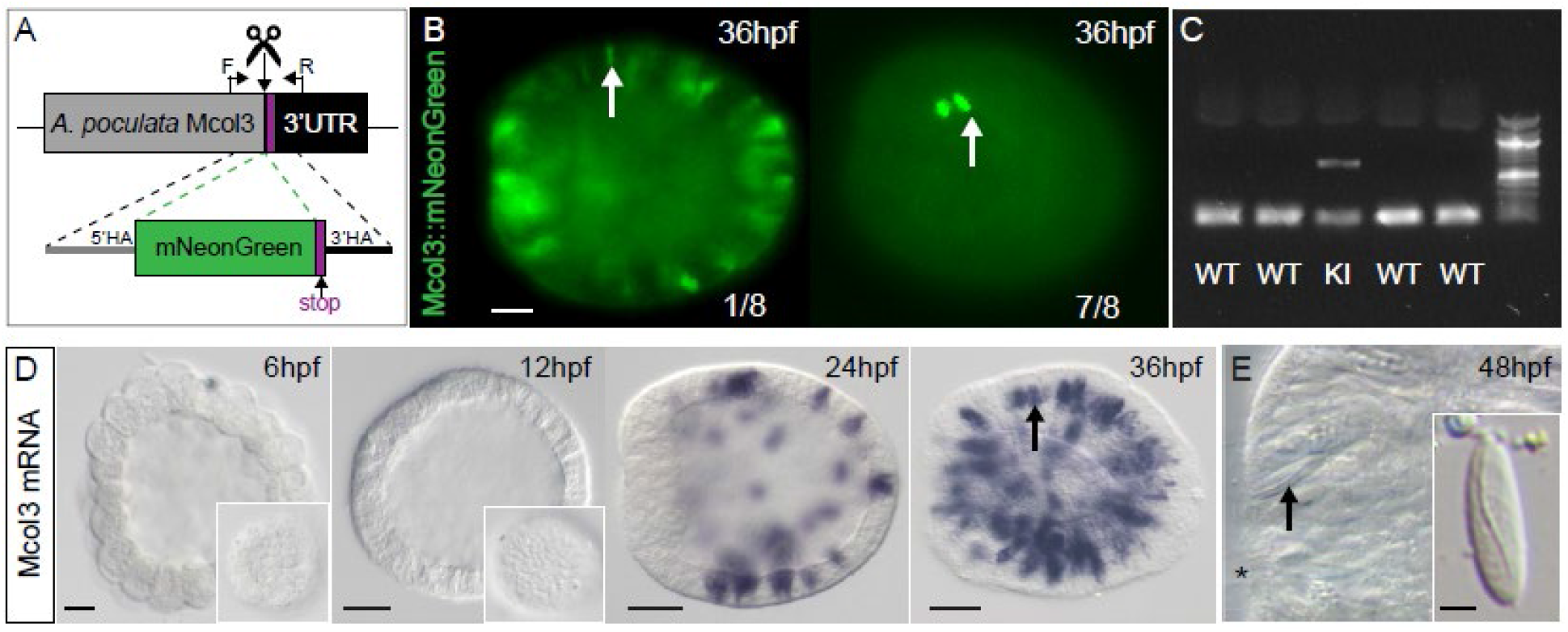
Endogenous labeling of developing cnidocytes using CRISPR/Cas9 genome editing. (A) Schematic showing knock-in strategy with relative position of sgRNA (scissors), genotyping primers (F/R, bent arrows), and repair template, including left and right homology arms (5’HA, 3’HA). The stop codon is indicated in purple. (B) Live images of embryos exhibiting either complete (1/8 embryos) or mosaic (7/8 embryos) fluorescent expression of *Mcol3*::mNeon; labeled cnidocytes (arrows) are distributed throughout the ectoderm. (C) Agarose gel genotyping of five individual embryos, one knock-in mutant (KI) and four wild types (WT). The wild type amplicon (350 bp) is present in all five embryos and the amplicon containing the mNeon insert (1060 bp) is present only in the mutant. (D) *In situ* hybridization confirms the timing and distribution of cells expressing *Mcol3* mRNA (immature cnidocytes) in the ectoderm at/after 24 hpf. Insets show surface detail and arrow points to *Mcol3*-expressing immature cnidocytes. (E) Mature cnidocytes are detected in the ectoderm at 48 hpf. DIC image of the oral region of a 48 hpf larva. Inset shows an isolated cnidocyte extracted from a dissociated larva; arrow points to a mature cnidocyte in situ. The oral pole is to the left in B,D-E; the position of the blastopore is marked by * in E. Scale bars in B,D = 20μm; scale bar in E = 2μm.

Mature cnidocytes are visible in the larva at 48 hpf, shortly after the onset of expression of *Mcol3* (**Fig 3E**). Together, these data show that the timing and distribution of fluorescent cells observed in knock-in larvae are consistent with the endogenous expression of *Mcol3* mRNA in *A. poculata* and the appearance of mature cnidocytes in wild type larvae. Cnidocytes are thought to have evolved from a neural-like precursor in the ancestor of cnidarians (Babonis et al., 2022), yet our understanding of the complex regulatory interactions that drive diversification of cnidocyte form and function remains limited (Babonis et al., 2023). The ability to track early cnidocyte development in vivo using endogenously tagged proteins in *A. poculata* makes this animal a critical model for understanding diversification of this phylum-restricted cell type.

## Conclusions

Due to the ease of collection and the ability to control the timing of spawns in the lab, *Astrangia poculata* is a tractable organism for functional genomic studies in hard corals. Their hardy, transparent embryos are robust to microinjection and genetic manipulation. We show that gene silencing and overexpression can be achieved by microinjection using low-cost techniques. We also show that exogenous gene knock-in can be readily achieved using a repair template generated by PCR to induce HDR following a CRISPR/Cas9 cutting reaction. The toolset presented here enables future studies of development in *A. poculata*, which has recently been recognized as a valuable experimental system for investigative studies of coral-microbe studies (Puntin et al., 2022), as a tractable research organism for functional studies in corals and, more broadly, as a viable system for comparative studies of cnidarian evolution and development (**Fig 4**). Additionally, we show that *A. poculata* is a valuable organism for studies of comparative developmental biology in cnidarians as this animal shares some larval features in common with *N. vectensis* and shares other features in common with other corals. We anticipate that the functional genomic techniques described here can be readily adapted for studying early development in other coral species and will accelerate research on fundamental cellular and molecular processes in corals and enable finer scale comparisons of comparative development in Anthozoa.

**Fig. 4.**
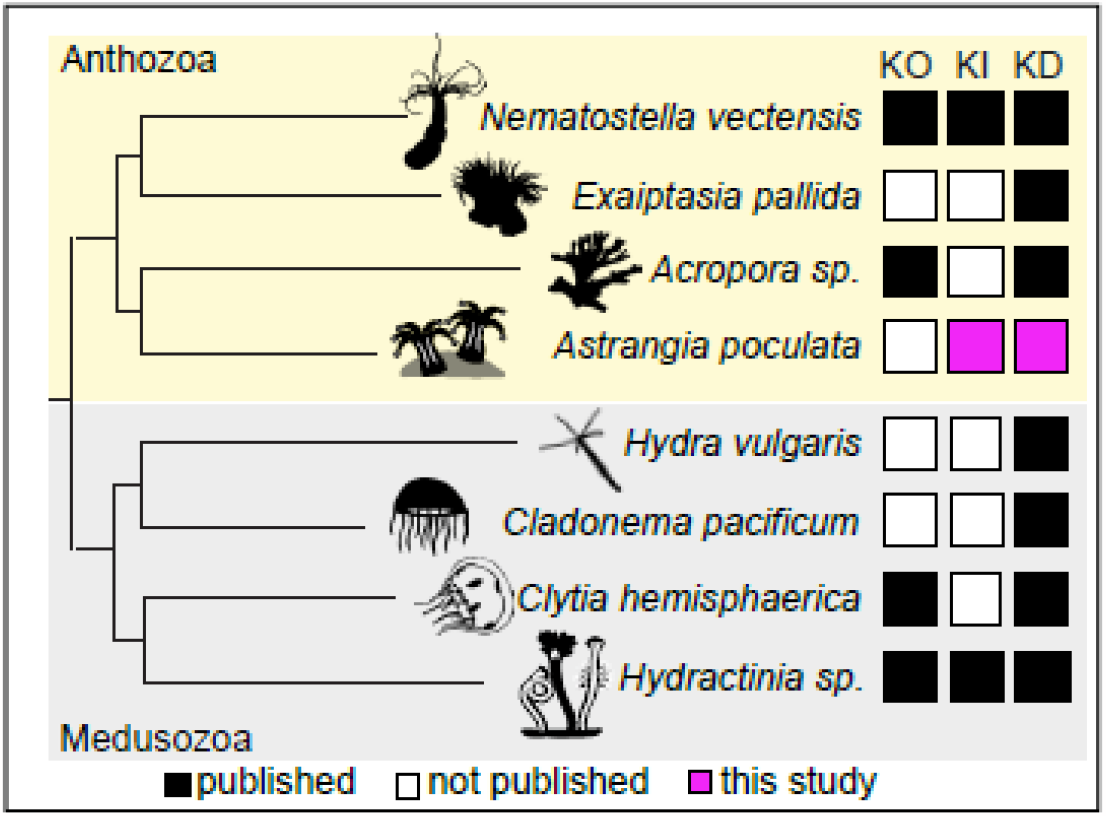
Summary of functional genomic tools available in cnidarians. KO – knockout, KI – knock-in, KD – knockdown by RNA interference. References by taxon: *Nematostella vectensis* (He et al., 2018; Ikmi et al., 2014), *Exaiptasia pallida* (Dunn et al., 2007), *Acropora millepora* (KO) (Cleves et al., 2018), *Acropora tenuis* (KD) (Yuyama et al., 2021), *Astrangia poculata* (this study), *Hydra vulgaris* (Lohmann et al., 1999), *Cladonema pacificum* (Masuda-Ozawa et al., 2022), *Clytia hemisphaerica* (KO) (Momose et al., 2018; Quiroga Artigas et al., 2018), *Clytia hemisphaerica* (KD) (Masuda-Ozawa et al., 2022), *Hydractinia echinata* (KO) (Gahan et al., 2017), *Hydractinia symbiolongicarpus* (KI,KD) (DuBuc et al., 2020; Quiroga-Artigas et al., 2020; Sanders et al., 2018). The cladogram was inferred from two studies of overlapping taxa (Fang et al., 2022; Kayal et al., 2018). Silhouettes were downloaded from Phylopic.org, license (CC BY-SA 3.0).

## Acknowledgements

This work was supported by the North Carolina Biotechnology Center (2022-FLG-3803, J.F.W.) and the National Institutes of Health (R15GM139113-01A1 to J.F.W., R35GM147253-01 to L.S.B., and P20GM103430 to K.S.)

## Author Contributions

JFW, RB, AS, EB, IVC, KS and LSB collected data and performed analyses; JFW and LSB conceived of the study and wrote the manuscript; RB, AS, EB, IVC, and KS edited and approved the manuscript.

## Data and Code Availability

The transcriptome assembly used for cloning, primer design, and knock-in construct design has been deposited at the NCBI repository (PRJNA956119) and is publicly available as of the date of publication. All transcript, primer, and donor sequences associated with this manuscript are provided in the Methods.

## Materials and Methods

### Animal collection and maintenance

Fresh colonies of *A. poculata* were collected by divers from Ft. Wetherill, RI, USA and transferred to Roger Williams University. Spawning was induced by acute heat shock (27-28°C for 1 hour) in benchtop containers. After spawning, adult colonies were maintained in a flow-through system with natural seawater and a 12:12 light cycle at Roger Williams University, Bristol, RI.

### Microinjection

*Astrangia poculata* gametes were collected and fertilized in 0.2um-filtered sea water and then transferred in filtered seawater to a 35mm petri dish containing 100μm mesh (Sefar Nitex 03-100/32) secured to the bottom using modeling clay. Individual zygotes were injected using a fluorescent Zeiss Discovery V8 dissecting scope, Narishige micromanipulator, and Eppendorf FemtoJet 4i picospritzing device, following a protocol developed previously for *N. vectensis* (Layden et al., 2013). Two different dextran dyes (Alexa 555 and Alexa 488 - Invitrogen D34679, D22910) each diluted to a final concentration of 0.2mg/ml in nuclease free water (Ambion AM9937) were used to mark individual blastomeres by injection at the two-cell stage. To assess the feasibility of expressing heterologous mRNA in *A. poculata*, zygotes (one-cell stage) were injected with mRNA encoding a NvTCF-venus fusion protein construct (Röttinger et al., 2012) diluted to 300ng/ul with 0.2mg/ml RNAse-free dextran in nuclease free water. Injected embryos were reared at room temperature (∼ 22°C) for both experiments, mounted in filtered sea water on glass slides, and imaged live on a Nikon Eclipse E800 fluorescent microscope at Roger Williams University.

### Transcriptome assembly

Larvae were collected at 12 hpf, 24 hpf, 36 hpf, 60 hpf, and 84 hpf in Tri-reagent (Sigma T9424) and stored at -80°C prior to RNA extraction. Total RNA was purified following a protocol previously described for *N. vectensis* (Layden et al., 2013). Briefly, samples were processed through two phenol/chloroform extractions and precipitated in isopropanol before being treated for DNA contamination with Turbo-DNAse (Ambion AM1907) for 10 in at 37°C. Library preparation and 150 bp PE illumina sequencing (NovaSeq 6000) was carried out by Novogene. Sequencing reads were combined and error corrected using Rcorrector (Song and Florea, 2015). Adapter trimming and quality trimming were carried out using Cutadapt v3.7 (Martin, 2011) and Trimmomatic v.039 (Bolger et al., 2014), respectively.

Cleaned reads were filtered for ribosomal sequences by aligning them ribosomal sequences for *A. poculata* from the SILVA database (Quast et al., 2013) using Bowtie2 v2.3.4.1 (Langmead and Salzberg, 2012). Unaligned reads were input into Trinity v2.12.0 (Grabherr et al., 2011) for assembly and final, assembled transcripts were filtered for sequences longer than 200 bp. Raw sequencing reads and assembled transcripts have been deposited to NCBI under bioproject: PRJNA956119.

### shRNA design and synthesis

The *A. poculata* ortholog of FgfA1 was identified using TBLASTN with the *N. vectensis* FGFA1 peptide sequence (NCBI accession: ABN70831.1) as query and the *A. poculata* assembled transcripts as reference. shRNAs were designed and synthesized as described previously for *N. vectensis* (He et al., 2018). In brief, primers were designed to target the 3’ end of the FgfA1 coding sequence using the Invivogen siRNA Wizard (www.invivogen.com/sirnawizard/design.php) and annealed for 2 min at 98°C in a thermocycler to generate a template for in vitro transcription. Transcription was performed using the Lucigen Ampliscribe T7 Flash kit (ASF3257) for 5h at 37°C in a thermocycler, following the manufacturer’s instructions. Products were column-purified using the Zymo Direct-zol RNA Miniprep Kit (R2050), aliquoted, and frozen at -80 C until the day of microinjection. A scrambled control shRNA was synthesized at the same time using primers described previously (Karabulut et al., 2019). All shRNAs were injected into zygotes at a concentration of 800ng/ul with 0.2mg/ml RNAse-free dextran in nuclease-free water. Embryos were reared to 48 hpf at room temperature, mounted in filtered seawater on glass slides, and imaged live on a Nikon Eclipse E800 at Roger Williams University. Primer sequences for FgfA1 shRNA synthesis are:

~~~
Apoc_FgfA1_shRNA_F:
TAATACGACTCACTATAGACAACAGCCGCATGACATTTCAAGAGAATGTCATGCGGCTGTTGTCTT
Apoc_FgfA1_shRNA_R:
AAGACAACAGCCGCATGACATTCTCTTGAAATGTCATGCGGCTGTTGTCTATAGTGAGTCGTATTA
~~~

### Fgf inhibitor treatment

Beginning immediately after fertilization, zygotes were incubated in 0.1% DMSO in filtered seawater containing 20mM SU5402 (Sigma SML0443) for a final concentration of 20μM SU5402 or 0.1% DMSO in filtered sea water (control). Embryos were reared at room temperature and solutions were refreshed every 24 hours until embryos were collected and fixed for immunostaining (48 hpf).

### Immunostaining

*Astrangia poculata* larvae were fixed in 4% paraformaldehyde (PFA) in filtered seawater and washed four times in phosphate buffered saline with 0.1% Tween-20 (PTw) for five minutes each. Non-specific protein interactions were blocked in 10% normal goat serum (NGS) diluted in PTw for 1 hour at room temperature. The blocking solution was replaced with a solution containing 1:200 anti-acetylated tubulin antibody (Sigma T6743) diluted in 10% NGS and the samples were incubated overnight at 4°C. Larvae were washed four times using PTw and incubated in a secondary antibody (Invitrogen A11004) diluted 1:200 in 10% NGS for 2 hours at room temperature. Larvae were again washed four times using PTw and counterstained in DAPI (Sigma D9542) diluted 1:2,500 and Phalloidin (Invitrogen A12379) diluted 1:200 in PBS overnight at 4°C. Larvae were washed four times in PTw, mounted in 75% glycerol in PBS on glass slides, and imaged on a Leica Sp8 confocal microscope at UNC Wilmington.

### In situ hybridization

Embryos from various developmental stages were collected and fixed for in situ hybridization (ISH) using a two-part fixative series. First, embryos were fixed for 1 min at room temperature in 4% PFA in PTw containing 0.25% gluteraldehyde. This initial fixative was removed and replaced with 4% PFA in PTw and embryos were fixed for an additional 1h at 4°C. Excess fixative was removed with three 10-min washes in PTw and tissues were then rinsed once in sterile water to remove excess PTw and twice in 100% methanol before being stored in clean 100% methanol at -20°C until analysis. ISH was performing following a method developed previously for *N. vectensis* (Wolenski et al., 2013), with minor modification. Due the small size and transparency of *A. poculata* embryos, all pipetting steps in the ISH procedure were performed in a sterile 24-well microplate on a dissecting microscope. An antisense mRNA probe directed against the *A. poculata Mcol3* transcript was synthesized as described for *N. vectensis* (Wolenski et al., 2013) using the following primers:

~~~
Apoc_Mcol3_F:
ATGGCGTCTAAACTCATTCTTG
Apoc_Mcol3_R:
TCACGCGTGCACACACCTA
~~~

Tissues were hybridized overnight with the Mcol3 probe diluted to 1ng/ul in hybridization buffer (Wolenski et al., 2013) and signal was visualized using an NBT/BCIP reaction performed in the dark at room temperature. Labeled embryos were washed extensively in PTw to remove excess NBT/BCIP, mounted in 80% glycerol (in PBS) on glass slides, and imaged on a Nikon Eclipse E800 at Cornell University.

### CRISPR-mediated knock-in

The *A. poculata* ortholog of *Mcol3* was identified using TBLASTN with the *N. vectensis* MCOL3 peptide sequence (NCBI accession: XP_032218917.1; Uniprot accession: G7H7X1) as query and the *A. poculata* assembled transcripts as reference. The open reading frame was predicted using the NCBI Open Reading Frame Finder (https://www.ncbi.nlm.nih.gov/orffinder/) and sgRNAs targeting the C-terminus of the predicted peptide were designed using ChopChop v3 (Labun et al., 2019). Two overlapping guides were designed with the recognition sites CGTGGTCGCTTACTTTCTGC **and** AATGTCGACGCATCATCACG that cut 4bp upstream and 11bp downstream from the insertion site respectively. Single guide RNAs (sgRNAs) were synthesized by Synthego (Redwood City, CA, USA) with the default 2’-O-Methyl modification at the 3 first bases and 3’ phosphorothioate bonds between the first three and last two bases.

Knock-in repair templates were synthesized using PCR. To do this, primers were designed to contain 40 bp homology arms that are homologous to the insertion site (immediately 5’ and 3’ to the predicted stop codon), a two-alanine spacer, and 15 bp to bind and amplify mNeonGreen in frame with the open reading frame. Silent mutations were introduced in the sgRNA recognition sequences to prevent recutting from sgRNAs. The oligo sequences were as follows (*Apoc* homology sequences, <u>sgRNA mutations</u>, [linker], **mNeon priming region**):

~~~
Apoc_Mcol3_Homology_F:
5’GCGTCTCGTCCTGCCCGACCCAGTGCTGCTCCGG<u>G</u>AG<u>G</u>AA<u>A</u>[GCCGCA]**ATGGTGAGCAAGGGC**3’
Apoc_Mcol3_Homology_R:
5’TAATTTCTAAATCTCGTGCTAATGTCGACGCATCA<u>AGTGC</u>T<u>C</u>GT<u>G</u>GC**TTACTTGTACAGCTCGTC**3’
~~~

PCR amplification of the repair template was performed using 50 ng of plasmid containing mNeonGreen (Addgene 125134) as template in a touchdown PCR reaction (annealing temperature decreased 1°C from 65°C to 50°C for the first 15 cycles followed by 20 cycles with an annealing temperature of 50°C; extension time was 30s). Afterwards, the template was digested by addition of DpnI enzyme (NEB R0176S) and incubation at 37°C for 2 hours. The repair template was then purified using a QiaQuick PCR purification kit (Qiagen 28704) prior to injection and quality assessed using agarose gel electrophoresis, to ensure size, and Nanodrop, to assess purity and concentration.

The injection mix was assembled as follows:

150ng/ul final concentration of dsDNA repair template

200ng/ul final concentration of ApMcol3 sgRNA1

200ng/ul final concentration of ApMcol3 sgRNA2

0.5ul of Cas9 protein (IDT 1081060)

Alexa-555 dextran (0.2mg/ml) Nuclease free water to 5ul

Ribonucleoprotein (RNP) assembly was promoted by incubation of the injection mix at room temperature for 15 minutes prior to injection.

### Imaging and genotyping knock-in mutants

Mutant embryos were identified using a fluorescent dissecting scope, mounted on glass slides in filtered sea water, and imaged live on a Nikon Eclipse E800 fluorescent microscope at Roger Williams University. Selected mutant knock-in larvae (48 hpf) were transferred individually to 0.5ml PCR tubes and gDNA was extracted from individual larvae as previously described (Servetnick et al., 2017). PCR was used to amplify the knock-in locus and insert size was confirmed using gel electrophoresis. Genotyping primers are as follows:

~~~
Apoc_Mcol3NG_F:
GGACCATCTGGACGAATGGGAC
Apoc_Mcol3NG_R:
CAATTCGCTCTTCTCTGCCTTCTAT
~~~

### Predicted gene sequences

*Astrangia poculata* Minicollagen 3 (constructed from multiple overlapping assembled transcripts,

*= stop codon location):

~~~
ATGAAAGACTCAACGACTGTCGAAATACAGCCTTATTCAAGCACATTTCAAAGATCGCTAGCCTGGGATCCAGAGATGGCGTCTAAACTCATTCTTGGGTGCTTAGCACTCATGGTAGTGTCGACCTACGCCAGATCAACATACAAAAGAAGCGCTAACCCGTGTCCCCCGGGATGTCCCGGTAGTTGTGCGCCCTCGTGTGCGGTGTCTTGTTGTCTTCCTCCACCACCCGCTCCACCACCGCCCCCACCCCCACCCCCACCACCACCAGAGCCCGCTAAGCCCGGACCACCTGGACCATCTGGACGAATGGGACCACCCGGACCTGTCGGACCTATCGGACCCATGGGAGAGGCCGGACCACCTGGAATACCCGGACCCCAAGGACCTCCTGGACCTCCCGGAGAACCCGCTCCTCCACCACCACCACCCCCACCGTGCCCACCTGTCTGCGCCCACACATGCGTCTCGTCCTGCCCGACCCAGTGCTGCTCCGGCAGAAAGTAA*GCGACCACGTGATGATGCGTCGACATTAGCACGAGATTTAGAAATTACTCCAACTTTAGCGTTCGTAAAGTACTTTTTCAGTGGA
~~~

*Astrangia poculata* FGF1A (CDS extracted from transcript TRINITY_DN12668_c0_g1_i4):

~~~
ATGAATTCCATTCAACTGCTTTTCCTACTTCAACTCTTTTGCTTCACGGAGATAAACACTTCAGCTAAACCGTACAACGCAACCAAATCCCAGACTAAAGATGCCGCGAGAACTTCAAGAGGATCTATCTCATCATCCATGACCAGATACGAAAACGACAGAATCAGAAACCATTCCCGAAAAACATTTCTTTCCAAGAAGCAGAAATGGCCACAACCGTCCACGGAAAGTTCCTTGAAACGTGTCCGCAAAATAACAAAACGACCGACAGCTACAACGCACTGTAAAATATTCTGCCGCAGCGGTTATCATCTTCAAATCCTTCCCAGTGGTGCAGTGAGGGGGACGGTTGACCAGGGCAGCAAGTACGTGTTGTTTGAGATGCAGTCATTTGGCCCTAGTCTCGTCAGGCTGATGAGTACAGCGACGGGCAGGTACCTATCTATGAGAAGAGACGGGAGTCTTCGAGGGTTGCGTAGCCAAAGTAACCGGGACTCACTTTTCAAAGAGACACATGAACAGAACGCGTTTCACTCTTACGCGTCACACAGATATTACAGACAACAGCCGCATGACATGTTGGTTGGCATCAAGAGAAACGGACAAATAAAACGAGCCACTAAAACCTTGCATGGACAAACTGCTACGCAATTTCTTGTCATCAAATTTTAA
~~~

### Quantification and Statistical Analysis

Quantitative analysis of apical tuft morphology was performed by measuring the maximum length of the longest aboral cilium and the length of the body axis (mouth to apical tuft base) in individual embryos using the Measure Tool in Fiji V1.54b (Schindelin et al., 2012). All data were analyzed in the R statistical computing environment V4.2.1 (R Core Team, 2020).

